# Logistic tumor-population growth and ghost-points symmetry

**DOI:** 10.1101/2023.08.30.555578

**Authors:** Stefano Pasetto, Isha Harshe, Renee Brady-Nicholls, Robert. A. Gatenby, Heiko Enderling

## Abstract

The observed time evolution of a population is well approximated by a logistic function in many research fields, including oncology, ecology, chemistry, demography, economy, linguistics, and artificial neural networks. Initial growth is exponential at a constant rate and capped at a limit size, i.e., the carrying capacity. In mathematical oncology, the carrying capacity has been postulated to be co-evolving and thus patient-specific. As the relative tumor-over-carrying capacity ratio may be predictive and prognostic for tumor growth and treatment response dynamics, it is paramount to estimate it from limited clinical data.

We show that exploiting the logistic function’s rotation symmetry can help estimate the population’s growth rate and carry capacity from fewer data points than conventional regression approaches. We test this novel approach against a classic oncology database of logistic tumor growth, achieving a 30% to 40% reduction in the time necessary to correctly estimate the logistic growth rate and carrying capacity. Our results will improve tumor dynamics forecasting and augment the clinical decision-making process.

## 1. Introduction

Mathematical modeling in biology and oncology has a long history(Araujo and McElwain 2004; Gerlee 2013; Ghaffari Laleh et al. 2022; Benzekry et al. 2014; Zahid, Mohsin, et al. 2021). The simplest known empirical model for population growth, the Malthusian growth model(Malthus 1798), is defined by exponential growth where the population concentration temporal evolution function *n* = *n*(*t*) is proportional to the growth speed *n*′ ≡ *dn*(*t*)/*dt* first time derivative of *n*. With the growth rate *γ* > 0 (units 1/time) and some initial conditions (i.c.) *n*(*t*_0_) = *n*_0_ at an arbitrary non-negative time *t*_0_ that we consider, without loss of generality, as the origin of the time axis *t*_0_ = 0, the system is given by *n*′ = *γn*. Analogous consideration holds for *γ* < 0 (i.e., decaying function), without significant changes. Given limited resources or self-competition as it grows, we can consider the generalization classically referred to as logistic growth, presented firstly by P. Verhulst in 1838 (Quetelet 1838). Logistic growth curves refer to the family of defined-positive two time-constant parameters {*κ*, *γ*}-curves, with *κ* referred to as the carrying capacity of the system. These curves are described mathematically as

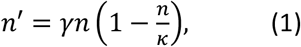

whose solution is fully algebraic for any given i.c. Using cancer as a demonstrative example, we will refer to *n*(*t*) as the clinically measurable tumor volume at the time, *t*. Of particular interest in the analysis of the logistic family of curves is the carrying capacity, *κ*, which represents the maximum population size that can be supported by the current conditions (such as the maximum tumor volume in an individual patient given the patient’s medical history (Hahnfeldt et al. 1999; Prokopiou et al. 2015; Sunassee et al. 2019; Poleszczuk et al. 2018; Zahid, Mohamed, et al. 2021)). To accurately predict the carrying capacity and the accompanying tumor growth dynamics, we may utilize the flex point *P*flex = *P*_flex_(*κ*, *γ*) of Eq.(1) reached at 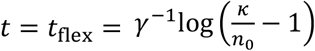 when 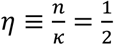, defined as the point at which the curvature changes sign (see Fig.1a). If any data set of disease dynamics follows logistic growth, the curve is entirely determined for *t* < *t*_flex_, as after *t*_flex_ the curve is rotationally symmetric with rotation θ = *kπ k* ∈ ℤ around *P*_flex_ (Fig. 1a).

**Figure 1.**
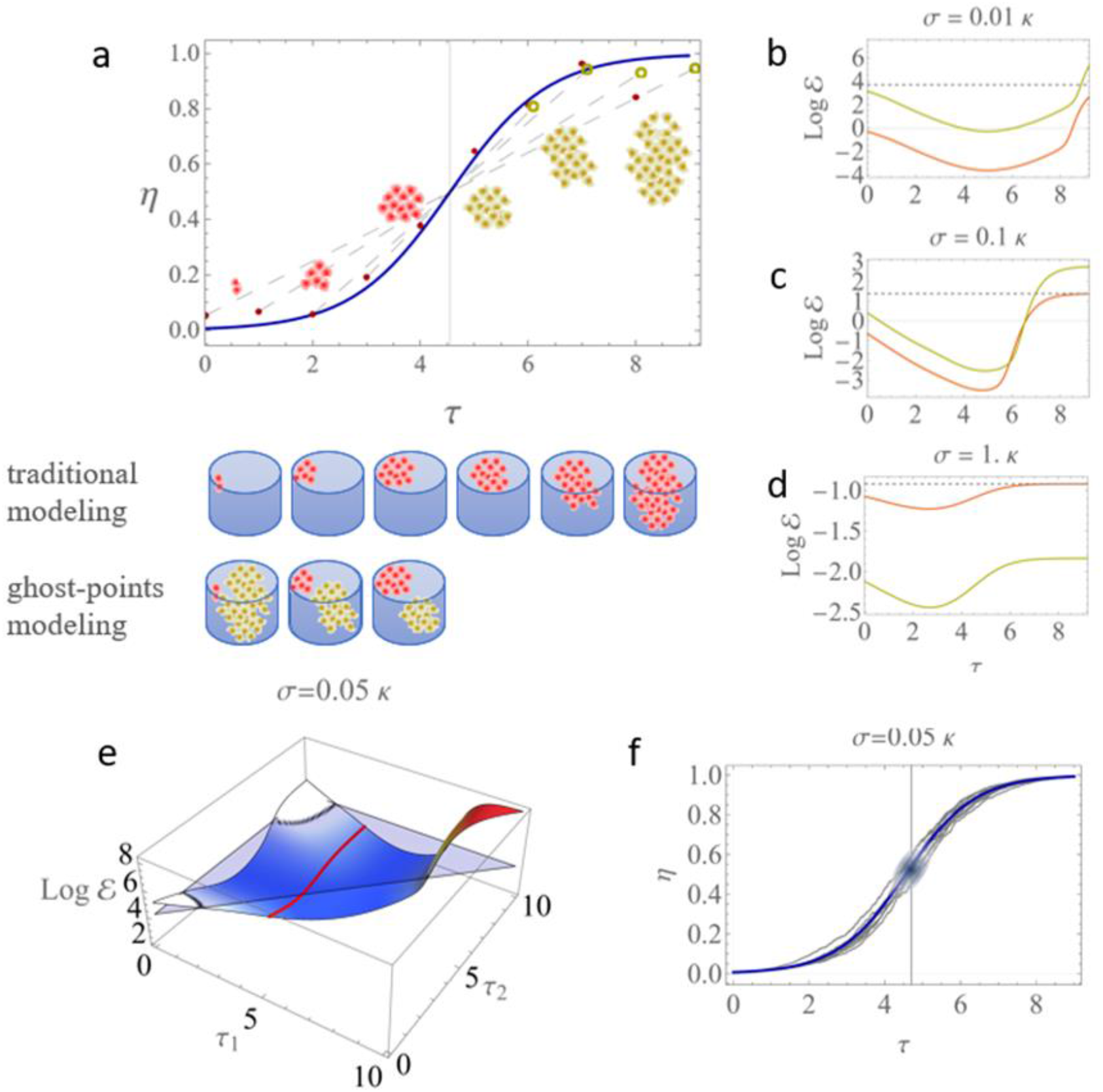
a) logistic trend and data collection with traditional (red cells) and ghost-point method (yellow cells). Non-dimensional quantities 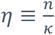 and 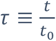 are drawn for aesthetic purposes, and dashed lines connect true to ghost points to guide the reader. In the Petri dishes pictorial representation, the ghost-points method allows us to obtain three guess values of the carrying capacity instead of waiting a long time (i.e., an earlier and more frequent determination). b, c, d) Logarithm of the evidence (eq. 2) of the classic (red) and ghost-point method (yellow) over the time evolution for three different data errors. The dotted lines correspond to a perfect detection (at null loglikelihood). e) time selection for a second follow-up tumor measurement: after a measurement at the time of diagnosis (τ= τ_1st_=4 in the figure) we can choose a second follow-up visit along the time axis of the second visit τ= τ_2nd_ by maximizing the evidence over the red line to exploit at the best the ghost-point method. f) error propagation in P_flex_ from stochastic analysis, have been tested (grey lines) to stay close to the correct profile (blue line) for σ < 0.1*κ*, see also panel c.

This study aims to demonstrate how *P*_flex_ can be used to predict how the tumor volume will evolve. We show how this simple consideration permits us to readily identify the logistic growth determination, i.e., the inference of the couple {*κ*, *γ*} of our cancer population with significant implications on oncological treatment selections (Zahid, Mohsin, et al. 2021).

## 2. Results

After the diagnosis of malignancy (say, *t*_0_ = 0), one or two follow-up measurements (say, point *P*_1_ at *n*_1_ and *t*_1_ and *P*_2_ at *n*_2_,*t*_2_) are usually performed at second opinion and eventual treatment planning. We can determine a “ghost-point” for each given data point, which is rotational-symmetric around *P*_flex_ (Fig. 1a). These ghost points permit us to estimate the logistic-growth asymptotic behavior far in the future and thus to have a predicted, more frequently sampled estimate of the determining couple {*κ*, *γ*}.

Let us call the ghost-point model *M*_*g*_, as compared to a classical regression model *M*_*c*_ where no ghost points are added. We want to understand under which condition *M*_*g*_ performs better than *M*_*c*_. It is intuitive to speculate that two factors play a significant role in this comparison: the moment the data points are collected and the error with which the observations are performed. The comparison between these two models of acquisition is best served within the Bayesian statistical framework, as it comes naturally equipped with dedicated mathematical instruments to compare the evidence ε of a model, say *M*_*g*_, over another, *M*_*c*_ (Stefano Pasetto, Gatenby, and Enderling 2021; S. Pasetto et al. 2021), by computing the evidence (or its log) traditionally as:

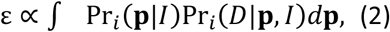

with *i* = {*g, c*}, **p** = {*κ*, *γ*}, and where the integral is extended to the entire parameter space of interest, representing the definition of model evidence apart from a constant normalization factor. The likelihood computation is different for the presence in Pr_*g*_(*D*∣p, *I*) of the ghost-point symmetry with respect to *P*_flex_ = *P*_flex_(*κ*, *γ*) (see Methods), where *D* represents the data collection *D* = {*P*_1_, *P*_2_, … } of real data points, *I* any other general information available (Stefano Pasetto, Gatenby, and Enderling 2021; Jaynes 2003). For example, carrying capacity can be characterized at the modeling level rather than as a free parameter, e.g., by imposing a max volume of stock inside a container, a max data transfer for fiber, the skull volume for brain cancer, etc., being these quantities sometimes referred as hyperparameters (i.e., parameters that do not enter in the inference problem). In medical terms, this information generally refers to as “anamnesis” of the patient. If a *I*= “patient is a smoker,” the prior information Pr_*i*_(p∣*I*) will encode a more discoraging prognosis in lung cancers (that can be parametrized via different shapes of Pr_*i*_(p∣*I*)). Similarly, if *I*= “the patient is a kid” the reduced skull volume size might be accounted in computing Pr_*i*_(*D*∣p, *I*) for brain cancers.

Fig. 1b-d shows that with sufficiently small errors, the ghost-point method has higher evidence and is consequently preferred compared to the classical regression model. In the figures, the data error σ is represented as a function of the carrying capacity *κ*; for example, in Fig1c, we suppose that each data carries an error bar of the order of 10% of the *κ*. This plot is informative of the importance at which each single data point is collected, as the evidence of *M*_*g*_ and *M*_*c*_ decreases of the same amount as time passes (*τ* in units the initial time *t*_0_, *τ* = *t*/*t*_0_), to growth again later on when the time approach the carrying capacity values. It is indeed well known experimentally that the detection time for a follow-up measurement of malignancy is relevant: early detection is preferable to a late one unless we are close to the maximum size permissible for the considered disease (i.e., the carrying capacity *κ*) assumed known (e.g., by body anatomical volume constraints). Nevertheless, the value of these comparative plots appears more apparent when they are read in relation one to the other, i.e., considering the vertical sequence of panels 1b to 1d of Fig. 1 carrying the growing error estimate over two orders of magnitude, from σ = 0.01*κ* to σ = *κ*. As the error increases, *M*_*c*_ overtakes *M*_*g*_ becoming the preferred model to use (Fig. 1d).

### 2.1 Implications for decision making

Of interest for mathematical oncology is the case of a second follow-up data point collection, say *P*_2_, as evidenced in Fig. 1e. We see how *M*_*g*_ augments the possibility of planning the best time for follow-up observations to predict disease evolution better as the model evidence ε not necessarily increases with time: If we follow the hypothetical clinical path of a patient in an oncological hospital, we can assume at *τ* = *τ*_*d*_, the time of the diagnosis, neoplasia becomes detectable in a patient by a diagnostic instrument (built with given precision σ = 0.05*κ*). We suppose that *τ*_*d*_ ≡ *τ*_1_ is the time of the first visit, see Fig. 1e, and it corresponds to a biomarker value, e.g., tumor volume, *η* = *η*_1_, whose expected growth is known to be logistic. We want to understand when the most convenient time for a follow-up visit *τ* = *τ*_2_ is (in order to use the ghost-points method optimally). From Fig. 1e, we follow the evolution of ε from *τ* = *τ*_1_ along with the red curve parallel to the *“τ*_2_*”*-axis. The evidence in favor *M*_*g*_ has a non-monotonic dependence on time *τ*: after an initial decrement, the evidence reaches a minimum and then grows up again (log ε are used in Fig. 1e in place of ε). Note how this is not always the case. If the patient reaches the diagnosis day “too late,” there is no advantage in a second follow-up measure (i.e., the red curve has crossed the plane of perfect detection, equivalent to the dotted lines in Figs 1b,c,d). This mathematical result states the importance of prevention and regular checkups in oncology. Following the same line of reasoning presented in Fig. 1e, we can imagine that clinical follow-up measurements, say *τ*_3_, *τ*_4_, and so forth, might require computational tools to be optimally established (a work beyond the goal of this paper). To conclude and recapitulate in a take-home message, in these series of figures, we stressed how, in a growing logistic process, the moment and the quality with which measurements are taken (i.e., when taken and with which precision) is more important than the number of measurements taken.

### 2.2 Patient-specific stochastic variation robustness

We test *M*_*g*_ against the ability to detect the *P*_flex_ = *P*_flex_(*κ*, *γ*) location. The flex point also depends on the parameters; therefore, checking the method’s robustness to its variability is essential. In particular, while *κ* is an extensive quantity (thus, it can be checked against literature, constrained with priors, or maximized by physical exams), *γ* is an intensive quantity. For this reason, while the knowledge of *κ* has extrinsic evidence in the nature of the system whose dynamics we want to study (e.g., of the order of mm^3^ or cm^3^ for tumors in a mouse or patient, respectively), a similar numerical rate *γ* can be reasonable for these very different systems (e.g., intrinsic cell proliferation happens on the order of once per day). In Fig. 1f, we check the robustness of *M*_*g*_ against stochastic variation to emulate patient sample heterogeneity. To test the robustness of the *M*_*g*_ against this critical dependency on *γ*, we assume fixed *κ*, and we proceed to the formal substitution *γ* → *γdt* + σ*dW* and *n*′ → *dn* into Eq. (1) with *dW* as the Wiener process, *γ* with a small abuse of notation is still-constant the growth rate, and σ → 0 reduces the Ito-process (i.e. the stochastic differential equation, SDE) to be deterministic. We integrate the corresponding SDE; a few examples are in Fig. 1f (Gallager 2013). *M*_*g*_ is consistently able to recover better values of {*κ*, *γ*} compared with *M*_*c*_ for the same range of errors studied in Figs 1b, c, d. See section Methods for technical details on determining the curves plotted in Fig1f.

### 2.3 Model forecasting

Finally, we perform *M*_*g*_ comparison against *M*_*c*_ on a truly observed dataset. For the sake of this work, we selected a well-known and standard dataset compiled by Laird (Laird 1964). The data map the growth for 19 samples of 12 different tumors of the rats, mice, and rabbits superimposed by scaling them according to the inflection point of the growth curve (see (Laird 1964) for further details on the original compilation). Because the trend of this compilation is logistic but involves different normalization scales as diverse as the animal size involved, the units of carrying capacity are normalized to a scale value *κ*, while the units of the growth rate are comparable, and we choose [*γ*] = [day^−1^]. We ignore the solution of the inference problem, briefly called true-values and given by p_*T*_ = {*κ*, *γ*}_*T*_ = {0.97(6) ± 4,0.76(8) ± 1} with uniform errors for all data σ = 0.04*κ*_*T*_, and we mimic the data collection as time passes.

In Fig.2a, we see the progression of the inference parameter over time. This result is compared with the same process for the method *M*_*g*_ in Fig. 2b. The advantage of *M*_*g*_ is evident in Fig.2c and Fig.2d, where the relative progression rescaled to the actual values are shown: *M*_*g*_ is able to recover the correct parameter values using data obtained three days earlier than *M*_*c*_, with an achievement quality evolution reminiscent of the arguments presented on the model evidence shown in Fig.1b-d. Finally, Fig. 2e and f compare the *M*_*c*_ and *M*_*g*_ posterior predictive distribution at the time of the flex point. From this, we see that *M*_*g*_ can correctly anticipate the actual trend of the disease progression.

**Figure 2.**
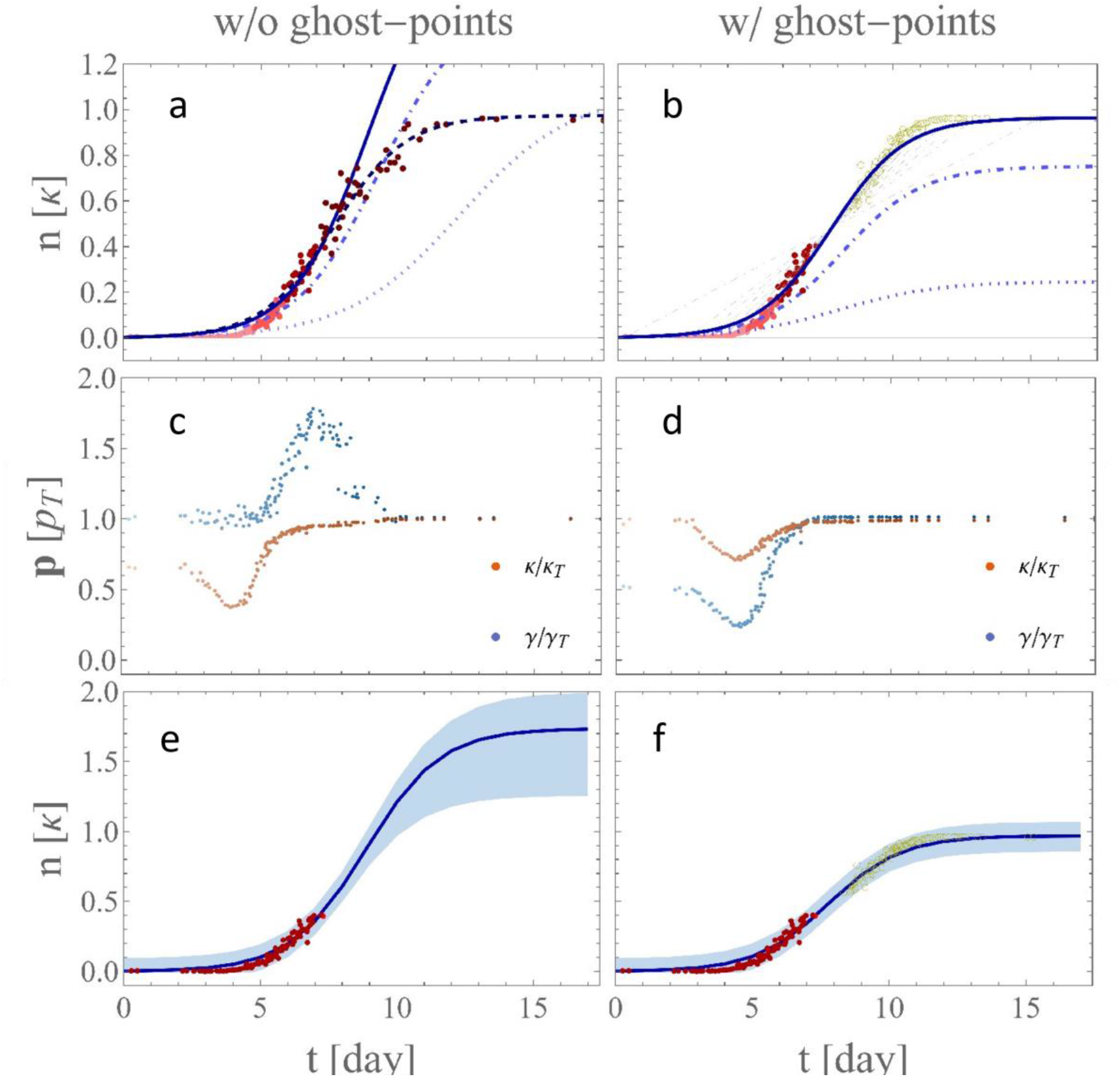
a and b) logistic curve inference (shown with dotted, dot-dashed, dashed, and solid lines) as data collection proceeds (corresponding red tonalities set-of-points from light to dark red) without and with the ghost-point method. C and d) parameter **p** = {*κ*, γ} normalized to true values without the ghost-point method. E and f) Bayesian posterior forecasting with the data collection at the flex point location. On panels b and f, yellow circles guide the reader by showing ghost points.

Finally, in Fig. 3, we present a specific case applicable to mathematical oncology. We consider triple-negative breast cancer priors informed by literature works (Nakashima et al. 2019). PDF of growing factor (Fig 3a) and carrying capacity (Fig 3b) are obtained from a beta prime distribution (Bourguignon 2021) fit to Table 2 of the Nakashima paper. Fig. 3c and e compare the forecasting bands at the time of the diagnosis and 50 days afterward without the help of ghost points for a chosen patient. Fig 3d and f present the same results with our novel technology. As evident, the patient-specific carrying capacity (around *κ*_*p*_∼2cm^*3*^) is reached earlier at parity of inspection times.

**Figure 3.**
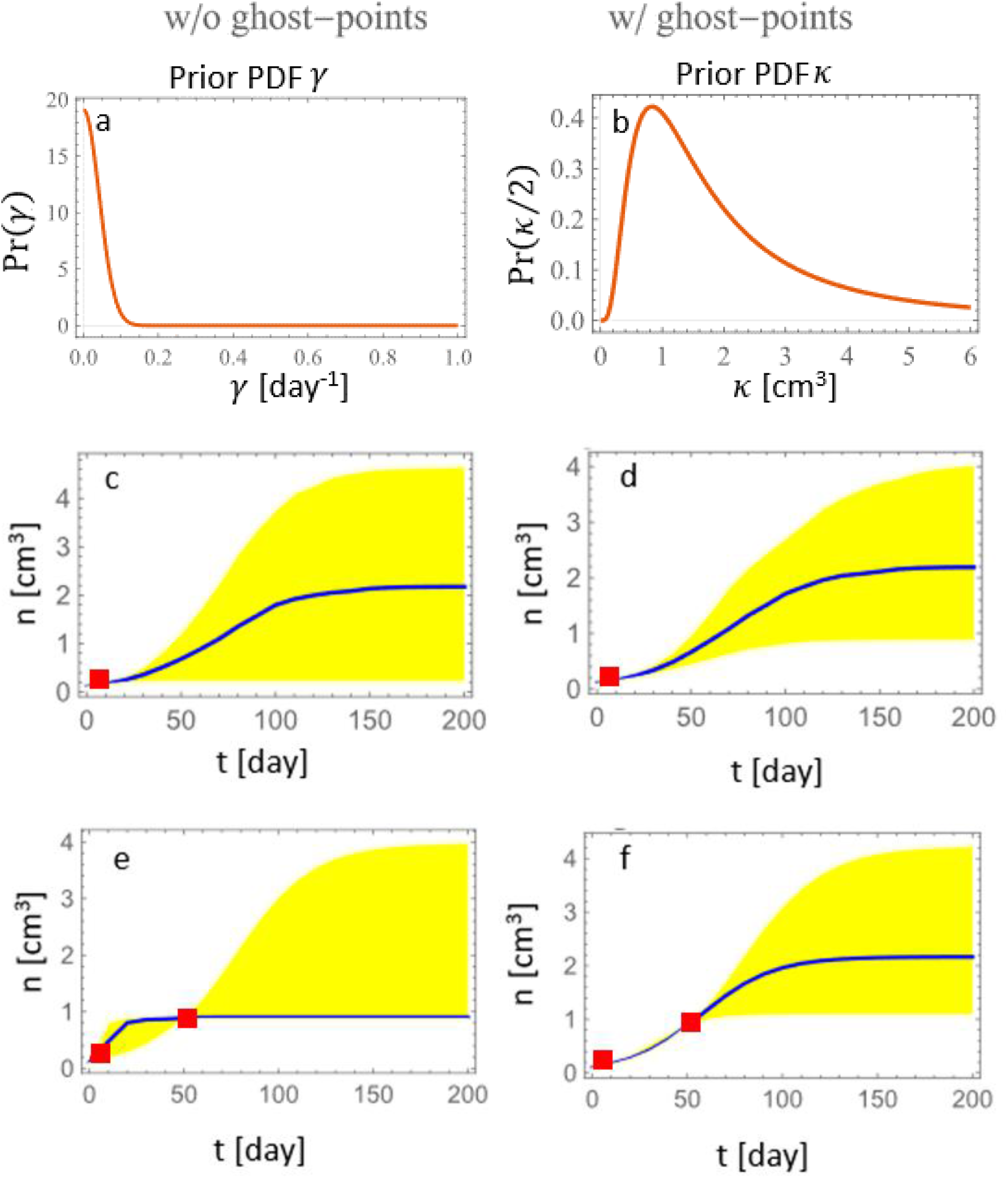
a) and b) prior probability distribution function for reaction rate and carrying capacity as from the dataset taken from the literature (see text). c) and d) prospected forecasted evolution at diagnosis computed as Figure 2. Yellow bands refer to the 5 and 90 percentile. e) and f) Bayesian posterior forecasting after the hypothetical second MRI/visit after 50 days. Error boxes on the collected data are shown in red.

## 3. Discussion

We focused on two crucial aspects of logistic-based analysis in medicine: to maximize information on tumor growth dynamics to optimize patients’ visit schedules and augment the logistic tumoral growth forecasting ability on true laboratory data. We demonstrated how a simple symmetry argument capitalizes the information acquired in the early stage of a disease with robust forecasting potentiality. This approach is expected to play a significant role in many research fields with logistic population dynamics. The critical dependence of the method on the quality of the data determination may be currently limiting, yet the utility of the ghost point symmetry approach is poised to increase with the continuous advancement of data collection technologies.

The introduced ghost point symmetry methodology has a comprehensive range of applications. Sigmoid functions (logistic, Gompertz, Richards’) are still the object of theoretical studies concerning problems of practical identifiability and data errors(Sharp et al., n.d.). Probably given the year of publication of the present work, the most remarkable application of logistics growth is related to the use of compartmental models in epidemiology. The SARS-CoV-2 virus that caused COVID-19 exhibited initial exponential growth flattened by implementing different countermeasures, including masking or physical distancing(Villalobos-Arias 2020; Postnikov 2020; Saito and Shigemoto 2020; Reiser 2021). Many applications that follow sigmoidal dynamics include the fields of Astrophysics(Kippenhahn and Weigert 1994) to map the probability distribution function of the electrons in degenerate white dwarfs stars, Chemistry(Yin and Zelenay 2018) to quantify the degradation of platinum group metal-free oxygen reduction reaction catalyst in fuel cell cathodes, Linguistics(“Probabilistic Linguistics” n.d.) to model language change, Economy(Rocha, Rocha, and Souza 2017) to track the public finance diffusion of credit pleas and the aggregate national response, and Oceanography to quantify the regrowth of coral(Simpson et al. 2022), to name but a few.

## 4. Methods

For the sake of completeness and more rapid reproducibility of the results, we present an explicit derivation of the loglikelihood part of Eq. (2), *L* ≡ Pr_*i*_(*D*∣p, *I*). For example, in the more elegant non-dimensional form, if we consider the general linear fractional transformation 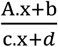 with affine matrix A ∈ *ℳ*(2,2), affine vector b ∈ *ℳ*(*m*, 1), fractional vector c ∈ *ℳ*(1, *n*) and fractional constant *d* ∈ *ℳ*(1,1) and where *ℳ*_*m*×*n*_ℝ the set of m-by-n real matrices, the staring (affine) transformation reads as:

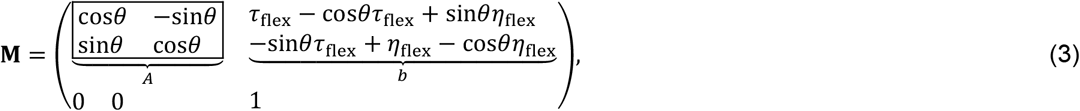

that applies to a generic point **p** = {*τ*_*p*_, *η*_*p*_}as

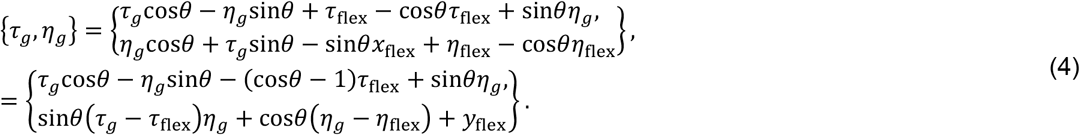

We then apply this transformation to each point (*τ*_*i*_, *η*_*i*_) to obtain each ghost point and compute the error of the ghost-point model on the point (i.e., the general likelihood term,*err*_*i*_). The general term in the likelihood *L* might be written in the form

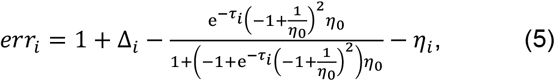

obtained by introducing the matrix-transformed general point in the logistic equation. We finally wrap it all in the loglikelihood rather than the simple likelihood to control numerical errors.Δ_*i*_∼*N*(**p**_*g*,*i*_, σ). The result is

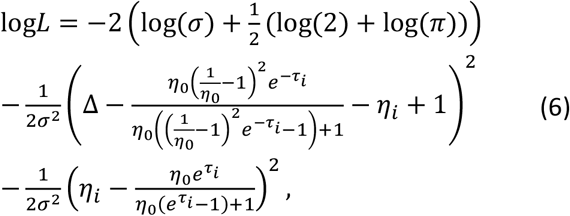

with the meaning of the symbols borrowed from the main text, summation over repeated index assumed, and Δ∼*N*(0, σ) randomizing the ghost-point location. Note that both Eq. (5) and (6) are further simplified, as well as the Gaussian perfector 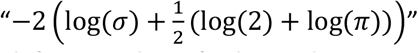 is always classically omitted in the computation. Here it is left uniquely to facilitate the equation derivation comprehension.

An introduction to the concept used but not developed in this work follows. This introduction is added upon request and not intended to be by any means comprehensive of all the technologies here exploited. Many books and original works are quoted in the reference where we point the reader interested in the original works together with white paper as Wikipedia, where most of these concepts are primarily explained.

### 4.1 Bayesian framework – a brief introduction

Equipped with the likelihood, several methodologies are available in the literature to solve the inference problem. Let’s refer with *d*_*i*_ to a generic data point from the dataset *D* with elements d = {*d*_1_, …, *d*_*n*_}, with p to the parameters of the data point distribution d∼Pr(d∣p), *I* all the extra information collectively referred to as hyperparameters *α* = {*α*_1_, …, *α*_*n*_}, i.e., p∼Pr(p∣*α*). Finally, with 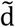 we refer to any new data point whose distribution is to be predicted.

The prior distribution, or Pr(p∣*I*), is the distribution of the parameter(s) before any data are observed. It may be difficult to determine the previous distribution; in this situation we just assume a poorly informative flat distribution. Furthermore, once equipped with the likelihood introduce in Eq. (6), from Bayes’ theorem results defined a posterior distribution as

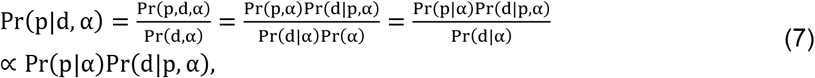

that represents also the integrand of Eq. (2). Finally, it remains define the posterior predictive distribution as the distribution of a new data point marginalized over the posterior:

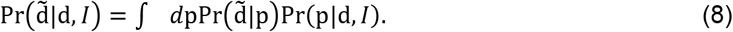

In order to do predictive inference, or to forecast the distribution of a brand-new, unobserved data point, Bayesian theory mandates the use of this posterior predictive distribution. In other words, a distribution of potential points is returned rather than a fixed point as a prediction. All the integrals means to be performed in ℝ^dim(p)^.

### 4.2 Nested sampling– a brief introduction

Numerical tacking the inference problem, as described above, has been for a long time a difficult challenge, surely not in our case with just a few parameters, but in the general case of an arbitrary number of parameters. Nevertheless, because we primarily aim at parameter inference and comparison between standard and ghost methods, a particularly successful approach can be used in our numerical solution of the Bayesian inference problem introduced above nested sampling. Nested Sampling (Skilling 2004) is a Monte Carlo algorithm, one of the most widely used Bayesian model comparison and inference techniques. It is a powerful technique for estimating the Bayesian evidence, which is a measure of the probability of the data under the model, and allows us to compare the likelihoods of different models too. The basic idea of nested sampling is to sample from the prior distribution while assigning each sample a weight proportional to the likelihood of the sample being the best sample within its level. A level is defined as a set of points with a fixed likelihood value. The algorithm starts by selecting an initial likelihood value and then finds the point with the smallest likelihood within that level. This point is considered to be the “worst” point in the level, and its likelihood is used to define the weight of the level. The algorithm then selects a new point with a likelihood smaller than the worst point in the previous level, and this process is repeated until the likelihood reaches a very small value. Once the algorithm has generated a set of samples, it uses them to estimate the Bayesian evidence as the sum of the products of the prior probabilities and the likelihoods of all the points in a given level. The final evidence estimate is obtained by summing the evidences of all levels. Finally, the algorithm generates samples from the posterior distribution, which is proportional to the product of the prior distribution (flat in our case) and the likelihood function (Eq. (6)), thus computing error bands in the forecasting problem. Here we outline a detailed implementation of our nested sampling algorithm stressing that this implementation is available in -virtually-any programming language:

a. We first define a prior distribution over the parameter space representative of the model. In our case, we can take p = {*n*_0_, *γ*, *κ*}. For our case a square box function has been assumed to modulate a flat distribution.
b. Generate *N*_*p*_ live points by sampling uniformly from the prior distribution, and calculate the likelihood *L*(θ) for each point with Eq (6). *N*_*p*_ ≥ 100 performs excellentlty on this database.
c. Set the initial “compression factor” *X* = 1.

We then perform the interactions:

1. we find the point with the lowest likelihood, *L*_mi*n*_, among the live points.
2. we estimate the evidence contribution from *L*_mi*n*_ as Δ*ε* = *L*_mi*n*_*X*/*N*_*p*_ and we accumulate this evidence by adding it to the running total *ε*_*tot*_ = *ε* + Δ*ε*.
3. We update the “dead” point with a new point sampled from the prior distribution but with a likelihood greater than*L*_mi*n*_. This new point replaces the dead point in the set of live points. 4) We update the compression factor: 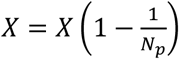 or, in our case, 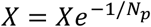 for better numerical stability.

Finally, we repeat the points 1 to 4 until a termination criterion is met, such as a target evidence accuracy, a maximum number of iterations, or a small change in the evidence. To finish, we calculate the remaining evidence contribution from the remaining live points and add it to the running total *ε*_*tot*_. The final estimate of the evidence, *ε* in Eq.(2) is given by the accumulated evidence,*ε*_*tot*_.

For the parameter estimation (the inference problem), we have that a) the posterior distribution can be approximated by the set of sampled points, their likelihoods, and their prior weights, and b) posterior samples can be obtained by resampling the sampled points with weights proportional to their likelihoods.

Generally, we note how the nested sampling algorithm is generally considered computationally intense, but it has several advantages over traditional methods like Markov Chain Monte Carlo (MCMC). These advantages include the ability to explore multi-modal distributions, better handling of degeneracies and strong correlations, and a more direct estimation of the evidence. Finally, note how several variants and improvements of the nested sampling algorithm have been developed, such as MultiNest (Feroz, Hobson, and Bridges 2009), PolyChord (Handley, Hobson, and Lasenby 2015), and Dynamic Nested Sampling (Higson et al. 2019) and many others, to address specific issues and improve the algorithm’s efficiency and accuracy in a many parameter setting (that we are not interested in considering here).

### 4.3 Stochastic differential equation – a brief introduction

Finally, *upon request of one of the referees*, we show the actual computation performed for the deterministic equation and the SDE mentioned above. Ito process, also known as an Ito-stochastic-process or a stochastic-differential-equation (SDE, as introduce above), are a mathematical family of models used to describe the behavior of random phenomena over time. An Ito process is a continuous-time stochastic process that evolves randomly over time, and its behavior is described by a differential equation that includes a stochastic component. In particular, the Ito process is defined by a stochastic differential equation of the form:

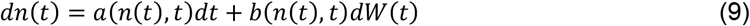

where *n*(*t*) is the process at time *t*, a(*n*(*t*), *t*) is the deterministic drift term, *b*(*n*(*t*), *t*) is the stochastic diffusion term, and *dW*(*t*) is a standard Wiener process (also known as Brownian motion). The stochastic differential equation describes how the process *n*(*t*) evolves randomly over time, where the drift term a(*n*(*t*), *t*) represents the expected change in *n*(*t*) over time, and the diffusion term *b*(*n*(*t*), *t*) represents the volatility or randomness of the process. The Wiener process *dW*(*t*) represents the random noise that drives the stochastic behavior of the process. Ito processes are widely used in finance, physics, and engineering to model systems that exhibit random behavior over time, such as stock prices, interest rates, particle motion, and diffusion processes.

For the simple goal of our project, we substitute the available algebraic solution of Eq. (1) with a numerical integration based on the Kloeden, Platen, and Schurz (KPS) (Kloeden, Platen, and Schurz 1994). KPS method is a standard numerical integration technique used to solve stochastic differential equations (SDEs). The KPS method is an extension of the well-known Euler-Maruyama method (Bayram, Partal, and Orucova Buyukoz 2018), a simple first-order numerical integration scheme for SDEs. KPS method belongs to a class of integration schemes called “strong order” methods, which provide accurate approximations of the underlying stochastic processes. The main idea behind the KPS method is to introduce higher-order terms in the Taylor series expansion of the solution to improve the accuracy of the approximation. Here is our brief outline of the KPS method for integrating the considered SDE:

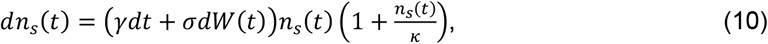

with *n*_*s*_ referring to the stochastic nature of this solution as compared to the deterministic one just named *n*. This equation clearly set in the general SDE of the form of Eq. (9). First, we proceed to discretize the time interval [0, *T*] into *N* equal subintervals, with a step size of *Δt* = *T*/*N*. For each time step, use the following approximation for the solution:

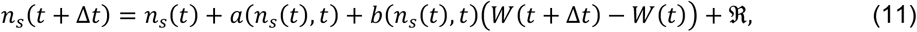

where ℛ is the remainder term that includes higher-order derivatives of a and *b*. The KPS method aims to improve the Euler-Maruyama method by including higher-order terms in the Taylor series expansion. The remainder term, ℛ, is given by:

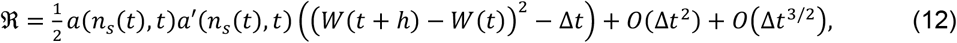

where a′(*n*_*s*_(*t*), *t*)denotes the derivative of the function a with respect to *n*_*s*_(*t*), and *O*(*Δt*^2^) and *O*(*Δt*^*3*/2^) represent higher-order terms that become smaller as the step size *Δt* decreases. To implement the KPS method, we perform the following steps for each time step from *t* = 0 to *t* = *T* − *Δt*:

a. Compute the increment in the Wiener process, Δ*W* = *W*(*t* + *Δt*) − *W*(*t*), a random variable is drawn from a normal distribution with mean 0 and variance *Δt*.
b. Calculate the deterministic and stochastic components of the SDE using *a*(*n*_*s*_(*t*), *t*) and *b*(*n*_*s*_(*t*), *t*).
c. Compute the higher-order term, ℛ, using Eq(12).
d. Update the state variable *n*(*t* + *Δt*) using the approximation formula:

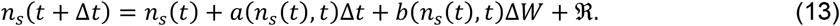

We then repeat steps a-d for each time step until the final time *T* is reached. The resulting sequence of *n*_*s*_(*t*) values provides an approximation of the solution to the SDE that substitute the algebraic *n*(*t*) of the non-stochastic (deterministic) available case. Note that the KPS method has a strong order of convergence equal to 1, which means that the error in the approximation decreases linearly as the step size *Δt* is reduced. This is an improvement over the Euler-Maruyama method, which has a strong order of convergence equal to 1/2.

### 4.4 Solution of the deterministic logistic differential equation

Finally, the implemented deterministic solution of Eq.(1) is achieved as follows. We separate the variables by writing Eq.(1) as

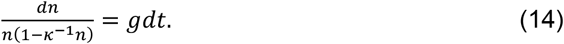

On the left-hand side (LHS), we use partial fraction decomposition to write

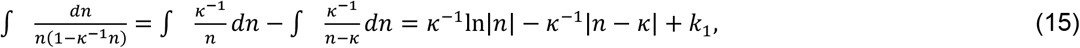

while the RHS gives *∫ gdt* = *gt* + *k*_2_, with integration constants *k*_*i*_. Finally, we use the initial condition to determine the integration constants. By substitution of *n*(*t*_0_) = *n*_0_ into the previous determined solution after simple algebraic manipulation the form we implemented:

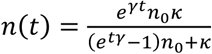

Note to the reader: while we validated the solution *n* = *n*(*t*) via algebraic manipulation, *we stress that we do not have taken any part in the realization or validation of KPS method for the computation of n*_*s*_ = *n*_*s*_(*t*), *we simply utilize numerical solvers that implement it (or any other equally valid method) and do not take the responsibility to validate or advertise any of them*.

## Acknowledgments

This work was supported by NIH/NCI grant U01CA244100; the JAYNE KOSKINAS TED GIOVANIS FOUNDATION FOR HEALTH AND POLICY, a private foundation committed to critical funding of cancer research. The opinions, findings, conclusions, or recommendations expressed in this material are those of the author(s) and not necessarily those of the JAYNE KOSKINAS TED GIOVANIS FOUNDATION FOR HEALTH AND POLICY, or their respective directors, officers, or staffs.

## Author Contributions

SP: conceptualized, developed, and realized the project. IH, RBN, RG, and HE contributed to the concept development exposition and funding collection.

## Data availability

The datasets used and/or analysed during the current study available from the corresponding author on reasonable request.

## References

Araujo, R. P., and D. L. S. McElwain. 2004. “A History of the Study of Solid Tumour Growth: The Contribution of Mathematical Modelling.” Bulletin of Mathematical Biology 66 (5): 1039. 10.1016/j.bulm.2003.11.002.

Benzekry, Sébastien, Clare Lamont, Afshin Beheshti, Amanda Tracz, John M. L. Ebos, Lynn Hlatky, and Philip Hahnfeldt. 2014. “Classical Mathematical Models for Description and Prediction of Experimental Tumor Growth.” PLoS Computational Biology 10 (8): e1003800. 10.1371/journal.pcbi.1003800.

Bourguignon, Marcelo. 2021. “A New Regression Model for Positive Random Variables with Skewed and Long Tail | SpringerLink.” 2021. https://link.springer.com/article/10.1007/s40300-021-00203-y.

Feroz, F., M. P. Hobson, and M. Bridges. 2009. “MultiNest: An Efficient and Robust Bayesian Inference Tool for Cosmology and Particle Physics.” Monthly Notices of the Royal Astronomical Society 398 (4): 1601–14. 10.1111/j.1365-2966.2009.14548.x.

Gallager, R.G. 2013. Stochastic Processes: Theory for Applications. Stochastic Processes: Theory for Applications. Cambridge University Press. https://books.google.com/books?id=ERLrAQAAQBAJ.

Gerlee, Philip. 2013. “The Model Muddle: In Search of Tumor Growth Laws.” Cancer Research 73 (8): 2407–11. 10.1158/0008-5472.CAN-12-4355.

Ghaffari Laleh Narmin, Chiara Maria Lavinia Loeffler, Julia Grajek, Kateřina Staňková, Alexander T. Pearson, Hannah Sophie Muti, Christian Trautwein, Heiko Enderling, Jan Poleszczuk, and Jakob Nikolas Kather. 2022. “Classical Mathematical Models for Prediction of Response to Chemotherapy and Immunotherapy.” PLoS Computational Biology 18 (2): e1009822. 10.1371/journal.pcbi.1009822.

Hahnfeldt, P., D. Panigrahy, J. Folkman, and L. Hlatky. 1999. “Tumor Development under Angiogenic Signaling: A Dynamical Theory of Tumor Growth, Treatment Response, and Postvascular Dormancy.” Cancer Research 59 (19): 4770–75.

Handley, W. J., M. P. Hobson, and A. N. Lasenby. 2015. “Polychord: Next-Generation Nested Sampling.” Monthly Notices of the Royal Astronomical Society 453 (4): 4384–98. 10.1093/mnras/stv1911.

Higson, Edward, Will Handley, Mike Hobson, and Anthony Lasenby. 2019. “Dynamic Nested Sampling: An Improved Algorithm for Parameter Estimation and Evidence Calculation.” Statistics and Computing 29 (September): 891–913. 10.1007/s11222-018-9844-0.

Jaynes, Edwin. 2003. Probability Theory: The Logic Of Science. Vol. 27. 10.1007/BF02985800.

Kippenhahn, R., and A. Weigert. 1994. Stellar Structure and Evolution.

Kloeden, Peter E., Eckhard Platen, and Henri Schurz. 1994. “Stochastic Differential Equations.” In Numerical Solution of SDE Through Computer Experiments, edited by Peter E. Kloeden, Eckhard Platen, and Henri Schurz, 63–90. Universitext. Berlin, Heidelberg: Springer. 10.1007/978-3-642-57913-4_2.

Laird, Anna Kane. 1964. “Dynamics of Tumour Growth.” British Journal of Cancer 18 (3): 490–502.

Malthus, Thomas. 1798. “An Essay on the Principle of Population.” In Wikipedia. https://en.wikipedia.org/w/index.php?title=An_Essay_on_the_Principle_of_Population &oldid=1007365843.

Nakashima, Kazuaki, Takayoshi Uematsu, Kaoru Takahashi, Seiichirou Nishimura, Yukiko Tadokoro, Tomomi Hayashi, and Takashi Sugino. 2019. “Does Breast Cancer Growth Rate Really Depend on Tumor Subtype? Measurement of Tumor Doubling Time Using Serial Ultrasonography between Diagnosis and Surgery.” Breast Cancer (Tokyo, Japan) 26 (2): 206–14. 10.1007/s12282-018-0914-0.

Pasetto, S., H. Enderling, R. A. Gatenby, and R. Brady-Nicholls. 2021. “Intermittent Hormone Therapy Models Analysis and Bayesian Model Comparison for Prostate Cancer.” Bulletin of Mathematical Biology 84 (1): 2. 10.1007/s11538-021-00953-w.

Pasetto, Stefano, Robert A. Gatenby, and Heiko Enderling. 2021. “Bayesian Framework to Augment Tumor Board Decision Making.” JCO Clinical Cancer Informatics, no. 5 (May): 508–17. 10.1200/CCI.20.00085.

Poleszczuk, Jan, Rachel Walker, Eduardo G. Moros, Kujtim Latifi, Jimmy J. Caudell, and Heiko Enderling. 2018. “Predicting Patient-Specific Radiotherapy Protocols Based on Mathematical Model Choice for Proliferation Saturation Index.” Bulletin of Mathematical Biology 80 (5): 1195–1206. 10.1007/s11538-017-0279-0.

Postnikov, Eugene B. 2020. “Estimation of COVID-19 Dynamics ‘on a Back-of-Envelope’: Does the Simplest SIR Model Provide Quantitative Parameters and Predictions?” Chaos, Solitons & Fractals 135 (June): 109841. 10.1016/j.chaos.2020.109841.

“Probabilistic Linguistics.” n.d. MIT Press (blog). Accessed October 17, 2022. https://mitpress.mit.edu/9780262523387/probabilistic-linguistics/.

Prokopiou, Sotiris, Eduardo G. Moros, Jan Poleszczuk, Jimmy Caudell, Javier F. Torres-Roca, Kujtim Latifi, Jae K. Lee, Robert Myerson, Louis B. Harrison, and Heiko Enderling. 2015. “A Proliferation Saturation Index to Predict Radiation Response and Personalize Radiotherapy Fractionation.” Radiation Oncology 10 (1): 159. 10.1186/s13014-015-0465-x.

Quetelet, Adolphe. 1838. Correspondance mathématique et physique. Impr. d’H. Vandekerckhove.

Reiser, Paul A. 2021. “Modified SIR Model Yielding a Logistic Solution.” arXiv. 10.48550/arXiv.2006.01550.

Rocha, Leno S., Frederico S. A. Rocha, and Thársis T. P. Souza. 2017. “Is the Public Sector of Your Country a Diffusion Borrower? Empirical Evidence from Brazil.” PLOS ONE 12 (10): e0185257. 10.1371/journal.pone.0185257.

Saito, Takesi, and Kazuyasu Shigemoto. 2020. “A Logistic Curve in the SIR Model and Its Application to Deaths by COVID-19 in Japan.” medRxiv. 10.1101/2020.06.25.20139865.

Sharp, Jesse A., Alexander P. Browning, Kevin Burrage, and Matthew J. Simpson. n.d. “Parameter Estimation and Uncertainty Quantification Using Information Geometry.” Journal of The Royal Society Interface 19 (189): 20210940. 10.1098/rsif.2021.0940.

Simpson, Matthew J., Alexander P. Browning, David J. Warne, Oliver J. Maclaren, and Ruth E. Baker. 2022. “Parameter Identifiability and Model Selection for Sigmoid Population Growth Models.” Journal of Theoretical Biology 535 (February): 110998. 10.1016/j.jtbi.2021.110998.

Skilling, John. 2004. “Nested Sampling.” AIP Conference Proceedings 735 (1): 395–405. 10.1063/1.1835238.

Sunassee, Enakshi D., Dean Tan, Nathan Ji, Renee Brady, Eduardo G. Moros, Jimmy J. Caudell, Slav Yartsev, and Heiko Enderling. 2019. “Proliferation Saturation Index in an Adaptive Bayesian Approach to Predict Patient-Specific Radiotherapy Responses.” International Journal of Radiation Biology 95 (10): 1421–26. 10.1080/09553002.2019.1589013.

Villalobos-Arias, Mario. 2020. “Using Generalized Logistics Regression to Forecast Population Infected by Covid-19.” arXiv. 10.48550/arXiv.2004.02406.

Yin, Xi, and Piotr Zelenay. 2018. “(Invited) Kinetic Models for the Degradation Mechanisms of PGM-Free ORR Catalysts.” ECS Transactions 85 (13): 1239. 10.1149/08513.1239ecst.

Zahid, Mohammad U., Abdallah S. R. Mohamed, Jimmy J. Caudell, Louis B. Harrison, Clifton D. Fuller, Eduardo G. Moros, and Heiko Enderling. 2021. “Dynamics-Adapted Radiotherapy Dose (DARD) for Head and Neck Cancer Radiotherapy Dose Personalization.” Journal of Personalized Medicine 11 (11): 1124. 10.3390/jpm11111124.

Zahid, Mohammad U., Nuverah Mohsin, Abdallah S. R. Mohamed, Jimmy J. Caudell, Louis B. Harrison, Clifton D. Fuller, Eduardo G. Moros, and Heiko Enderling. 2021. “Forecasting Individual Patient Response to Radiation Therapy in Head and Neck Cancer With a Dynamic Carrying Capacity Model.” International Journal of Radiation Oncology, Biology, Physics 111 (3): 693–704. 10.1016/j.ijrobp.2021.05.132.

